# Regulatory small RNA, Qrr2, is expressed independently of sigma factor-54 and functions autonomously in *Vibrio parahaemolyticus* to control quorum sensing

**DOI:** 10.1101/2021.07.01.450815

**Authors:** J.G. Tague, J. Hong, S.S. Kalburge, E.F. Boyd

## Abstract

Bacterial cells alter gene expression in response to changes in population density in a process called quorum sensing (QS). In *Vibrio harveyi*, LuxO, a low cell density activator of sigma factor-54 (RpoN), is required for transcription of five non-coding regulatory sRNAs, Qrr1-Qrr5, which each repress translation of the master QS regulator LuxR. *Vibrio parahaemolyticus*, the leading cause of bacterial seafood-borne gastroenteritis, also contains five Qrr sRNAs that control OpaR (the LuxR homolog), required for capsule polysaccharide (CPS) and biofilm production, motility, and metabolism. We show that in a Δ*luxO* deletion mutant, *opaR* was de-repressed and CPS and biofilm were produced. However, in a Δ*rpoN* mutant, *opaR* was repressed, no CPS was produced, and less biofilm production was observed compared to wild type. To determine why *opaR* was repressed, expression analysis in Δ*luxO* showed all five *qrr* genes were repressed, while in Δ*rpoN* the *qrr2* gene was significantly de-repressed. Reporter assays and mutant analysis showed Qrr2 sRNA can act autonomously to control OpaR. Bioinformatics analysis identified a sigma-70 (RpoD) -35 -10 promoter overlapping the canonical sigma-54 (RpoN) promoter in the *qrr2* regulatory region. Mutagenesis of the sigma-70 -10 promoter site in the Δ*rpoN* mutant background, resulted in repression of *qrr2.* Analysis of *qrr* quadruple deletion mutants, in which only a single *qrr* gene is present, showed that only Qrr2 sRNA can act autonomously to regulate *opaR*. Mutant and expression data also demonstrated that RpoN and the global regulator Fis act additively to repress *qrr2*. Our data has uncovered a new mechanism of *qrr* expression and shows that Qrr2 sRNA is sufficient for OpaR regulation.

**Importance:** The quorum sensing non-coding sRNAs are present in all *Vibrio* species but vary in number and regulatory roles among species. In the Harveyi clade, all species contain five *qrr* genes that, in *V. harveyi*, are additive in function to control LuxR. In the Cholerae clade, four *qrr* genes are present, and in *V. cholerae* the *qrr* genes are redundant in function to control HapR (the LuxR homolog). Here, we show that in *V. parahaemolyticus*, only *qrr2* can function autonomously to control OpaR, and it is controlled by two overlapping promoters. The *qrr2* sigma-70 promoter is present in all strains of *V. parahaemolyticus* and in other members of the Harveyi clade suggesting a conserved mechanism of regulation.

## Introduction

Bacteria monitor changes in cell density using a process termed quorum sensing (QS) (1, 2). QS is a regulatory mechanism used to alter global gene expression in response to cell density changes (1-6). In many Gram-negative bacteria, N-acylhomoserine lactone (AHL) is a common QS autoinducer synthesized intracellularly and secreted out of the cell (2, 7). By surveying AHL levels in its environment, a bacterium can regulate gene expression in response to growth phase. Quorum sensing has been characterized in several marine species in the genus *Vibrio*, including *V. anguillarum, V. cholerae, V. harveyi* and *V. parahaemolyticus*, and shown to modulate expression of bioluminescence, capsule formation, biofilm, natural competence, swarming motility, and virulence (5, 7-21). In *V. harveyi* and *V. anguillarum*, it was shown that LuxO, the QS response regulator, is an activator of sigma factor-54, encoded by *rpoN* that along with RNA polymerase, initiates transcription of the non-coding quorum regulatory small RNAs (Qrr) (8, 22-23).

Non-coding sRNAs are a group of regulators present in prokaryotes that together with the RNA chaperone Hfq control gene expression in a range of phenotypes (24-26). The Qrr sRNAs are classified as *trans*-acting sRNAs that along with Hfq, target mRNA via base-pairing to the 5′ UTR to stabilize or destabilize translation. In *V. harveyi,* the nucleoid structuring protein Fis was shown to be a positive regulator of *qrr* gene expression (27). The Qrr sRNAs are post-transcriptional regulators that, in *V. harveyi*, enhanced translation of the QS low cell density (LCD) master regulator AphA and inhibited translation of the QS high cell density (HCD) master regulator LuxR (28-31). At HCD in *V. harveyi*, LuxO is not phosphorylated and therefore cannot activate sigma-54 (RpoN), the five Qrr sRNA genes *qrr1* to *qrr5* are not transcribed, and LuxR translation is de-repressed. In addition, AphA and LuxR repress each other transcriptionally, providing a further level of regulation (30, 32-34). Studies have shown that in *V. harveyi*, Qrr1 has a 9-bp deletion in the 5’ region of the sRNA and therefore cannot activate *aphA* translation but can still repress *luxR* translation. The deletion in *qrr1* is also present in *V. cholerae, V. parahaemolyticus* and several other *Vibrio* species (30, 35). In *V. harveyi*, Qrr2, Qrr3, Qrr4, and Qrr5 sRNAs were additive in function and controlled the same target sites (29, 36, 37). However, the *qrr* genes showed distinct expression patterns and controlled the QS output signal at different levels coordinated with highest to lowest expression: Qrr4 > Qrr2 > Qrr3 > Qrr1 > Qrr5 (29). *V. cholerae* encodes four Qrr sRNAs, Qrr1 to Qrr4 that were redundant in function with any one of the four Qrr sRNAs sufficient to repress HapR (the LuxR homolog) (37).

*Vibrio parahaemolyticus* (VP) is a halophile, residing in marine environments as free-living organisms or in association with marine flora and fauna (38-40). This species is the leading cause of seafood-borne bacterial gastroenteritis worldwide, causing increasing infections each year, and is also a serious pathogen in the aquaculture industry (41, 42). *Vibrio parahaemolyticus* has dual flagellar systems, with the lateral flagellum system required for swarming motility, an important multicellular behavior (43). *Vibrio parahaemolyticus* has the same QS components and pathway as *V. harveyi*, containing five Qrr sRNAs that are predicted to control *aphA* and *luxR* (Fig. 1). In this species, the LuxR homolog is named OpaR (<u>Opa</u>city <u>R</u>egulator), for its role as an activator of capsule polysaccharide (CPS) production that results in an opaque, rugose colony morphology (44). A Δ*opaR* mutant has a translucent, smooth colony morphology and does not produce CPS nor a robust biofilm (14, 16, 44, 45). Besides CPS and biofilm formation, OpaR has also been shown to regulate swimming and swarming motility, surface sensing, metabolism, and the osmotic stress response in this species (14, 15, 20, 46-49). A *V. parahaemolyticus* Δ*luxO* deletion mutant, in which the *qrr* sRNAs were not expressed, showed *opaR* was highly induced and produced both CPS and biofilm, output signals of the QS pathway (14). Interestingly, an earlier study examining an Δ*rpoN* deletion mutant in *V. parahaemolyticus* showed that it did not produce CPS nor biofilm (50). This is unexpected because previous studies in *V. harveyi* suggest that in a Δ*rpoN* mutant the *qrr* sRNA would not be expressed and therefore *luxR* (*opaR*) would be de-repressed and production of CPS and biofilm would be observed.

**Figure 1:**
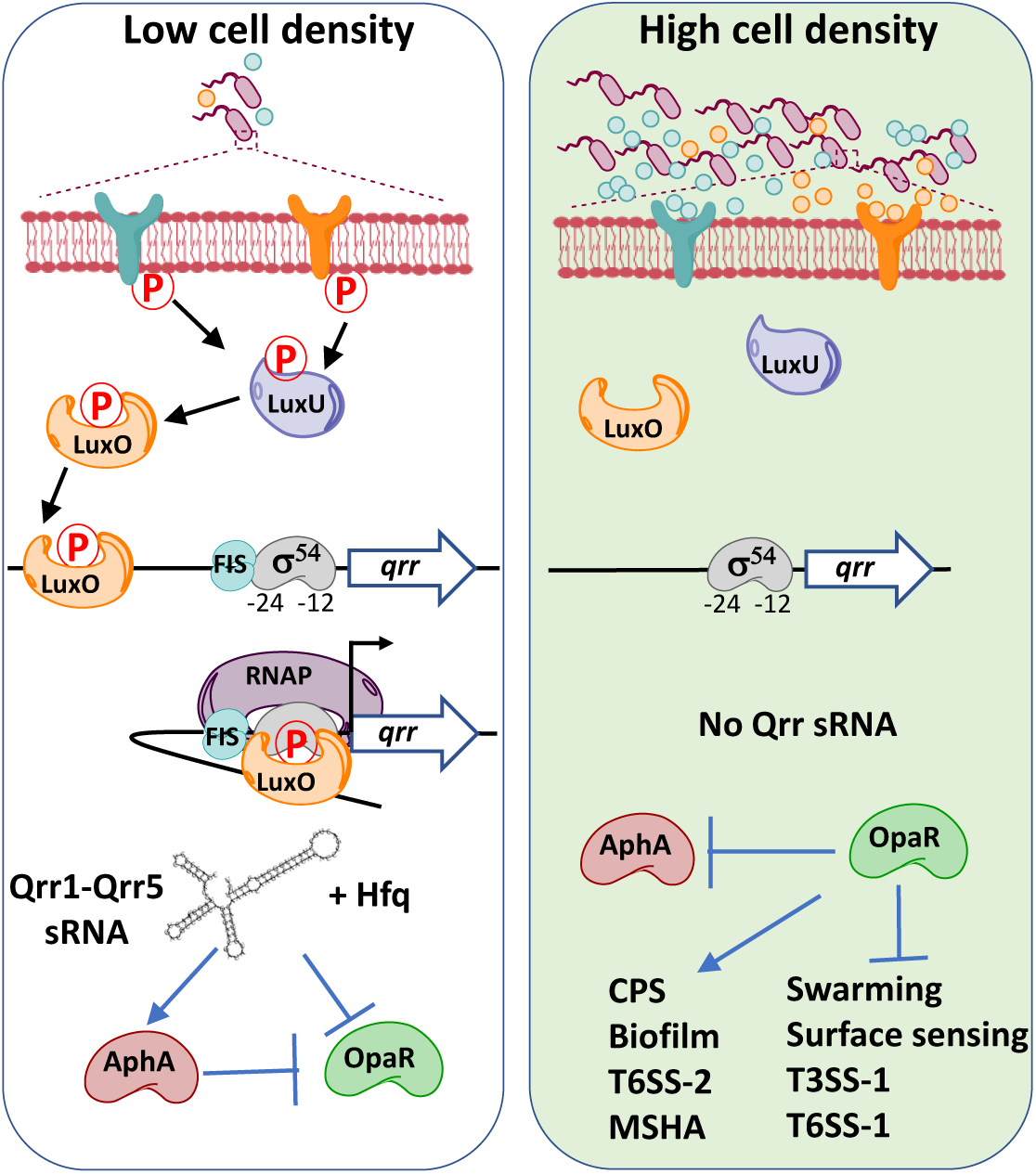
*Vibrio parahaemolyticus* quorum sensing pathway. Autoinducers (AIs) are synthesizes internally by three synthases and then excreted outside the cell. At low cell density, three histidine-kinase receptors are free of AIs, therefore act as kinases, phosphorylating LuxU and ultimately LuxO. LuxO-P activates RpoN and, along with Fis positively regulates transcription of five small quorum regulatory RNAs (Qrr sRNAs). The Qrr sRNAs, along with Hfq, stabilize *aphA* transcripts and destabilize *opaR* transcripts. In addition, AphA is a negative regulator of *opaR* expression. At high cell density, LuxO is unphosphorylated and inactivate, no *qrrs* are transcribed, *opaR* is expressed and *aphA* is repressed. OpaR positively regulates capsule polysaccharide production (CPS), biofilm formation, type 6 secretion system-1, and the type IV pilin MSHA, among other genes. OpaR negatively regulates swarming motility, surface sensing and two contact dependent secretion systems T3SS-1 and T6SS-1.

Here, we examined mutants of the QS pathway in *V. parahaemolyticus* to determine why the QS pathway output phenotypes differ between the Δ*luxO* and Δ*rpoN* mutants. We examined single and double mutants of Δ*luxO* and Δ*rpoN* for CPS and biofilm formation. We demonstrate that the Δ*luxO* mutant produces CPS and biofilm, whereas the Δ*rpoN* and Δ*rpoN/*Δ*luxO* mutants do not, suggesting *opaR* is repressed. We determined the expression patterns of *opaR, aphA* and the five *qrr* genes in these mutants and from this data, we determined that *qrr2* was de-repressed in the Δ*rpoN* mutant and *opaR* was repressed. Deletion of *qrr2* in the Δ*rpoN* mutant background resulted in *opaR* expression and restored CPS and biofilm production, demonstrating that Qrr2 sRNA is responsible for the Δ*rpoN* mutant CPS defect phenotype.

Bioinformatics analysis of the *qrr2* regulatory region identified an RpoD-35 -10 promoter region that overlaps with the RpoN -24 -12 promoter suggesting a mechanism by which *qrr2* is expressed in the *rpoN* mutant. We performed mutagenesis of the putative -10 promoter site and showed that *qrr2* expression was repressed indicting that *qrr2* can be transcribed from this promoter. To determine whether the other Qrr sRNAs can also function independently, we constructed quadruple *qrr* mutants, in which only one *qrr* is present, and examined CPS and motility phenotypes. Only the quad mutant containing *qrr2* could recapitulate wild type phenotypes, indicating that it can act autonomously to regulate OpaR. Furthermore, in a sigma-54 and Fis double mutant (Δ*rpoN/*Δ*fis*), *qrr2* was more highly expressed than in a single Δ*rpoN* mutant, suggesting that both RpoN and Fis act together to repress *qrr2* transcription by sigma-70. Sequence comparative analysis showed that -35 and -10 promoter sites were conserved among species within the Harveyi clade. Overall, our data show that Qrr2 can function independently and has a novel mechanism of expression in *V. parahaemolyticus*.

## Results

### Differential expression of *opaR* and *aphA* in Δ*luxO* versus Δ*rpoN* mutants

We used CPS production as a readout of OpaR presence in the *V. parahaemolyticus* cell. Based on the quorum sensing pathway in *V. harveyi*, we would expect both a Δ*luxO* and Δ*rpoN* deletion mutant to produce CPS, as the Qrr sRNAs should not be transcribed, and therefore OpaR should be de-repressed (**Fig. 1**). In a CPS assay, the Δ*luxO* mutant produced CPS forming opaque, rugose colonies. However, the Δ*rpoN* mutant did not produce CPS, instead forming a translucent, smooth colony morphology, similar to the Δ*opaR* strain (**Fig. 2A**). In addition, a Δ*rpoN/*Δ*luxO* double mutant also lacked CPS and produced a translucent, smooth colony morphology. Similarly, when we examined biofilm formation, both the Δ*rpoN* and Δ*rpoN/*Δ*luxO* double mutant strains produced less biofilm than the wild type and Δ*luxO* strains (**Fig. 2B**). These data suggest that in the Δ*rpoN* mutant, *opaR* is repressed. To test this, we complemented the Δ*rpoN* mutant with the *opaR* gene expressed from an arabinose promoter (pBAD*opaR*), and in these cells CPS production was restored, indicating that the absence of *opaR* in the Δ*rpoN* mutant led to the CPS defect (**Fig. S1**).

**Figure 2:**
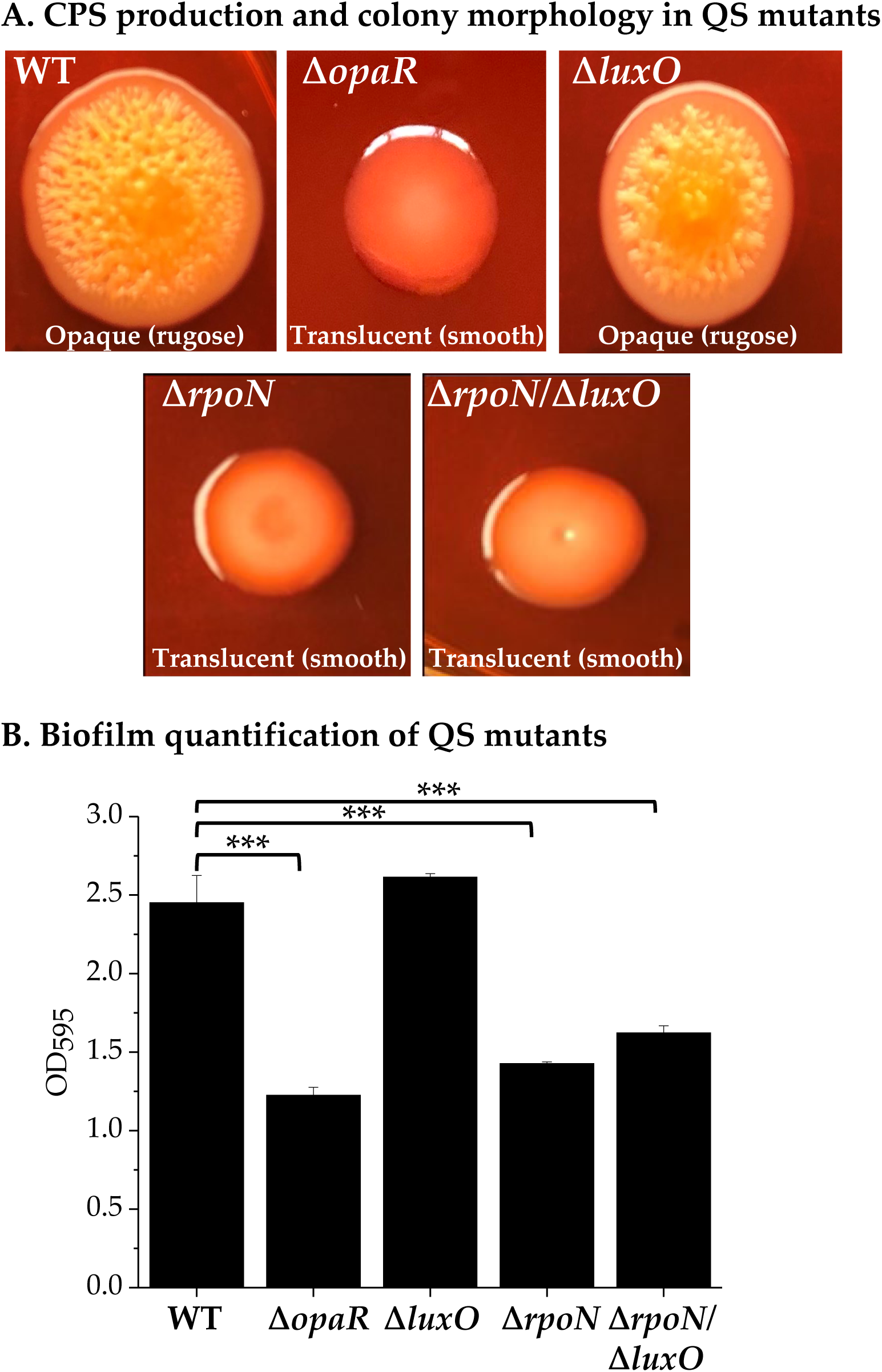
**A.** Wild type (WT) and QS mutant strains production of capsule polysaccharide (CPS) and colony morphology on Congo red plates. **B**. Biofilm assay from cultures grown for 24 h, stagnant and stained with crystal violet. Images are representatives from three bio-reps. Biofilm quantification of three bio-reps in duplicate. Statistics calculated using a Student’s t-test. ***, P-value <0.001

Next, we investigated the expression profiles of the QS master regulators in the Δ*luxO* and Δ*rpoN* deletion mutants. RNA isolation and quantitative real-time PCR (qPCR) assays were performed from cells grown in LB 3% NaCl to optical densities (OD) 0.1 and 0.5. At OD 0.1, expression of *opaR* in Δ*luxO* relative to wild type was significantly upregulated, however, *opaR* expression was unchanged in the Δ*rpoN* mutant (**Fig. 3A**). At OD 0.5, expression of *opaR* in Δ*luxO* matched that of wild type, however, in the Δ*rpoN* mutant expression of *opaR* was significantly downregulated relative to wild type (**Fig. 3B**). Expression of *aphA*, the low cell density QS master regulator, was repressed in the Δ*luxO* mutant compared to wild type and unchanged in the Δ*rpoN* mutant at OD 0.1 (**Fig. 3C**). At OD 0.5, *aphA* expression was upregulated compared to wild type in Δ*rpoN* (**Fig. 3D**). These data indicate that *opaR* is repressed in the Δ*rpoN* deletion mutant.

**Figure 3:**
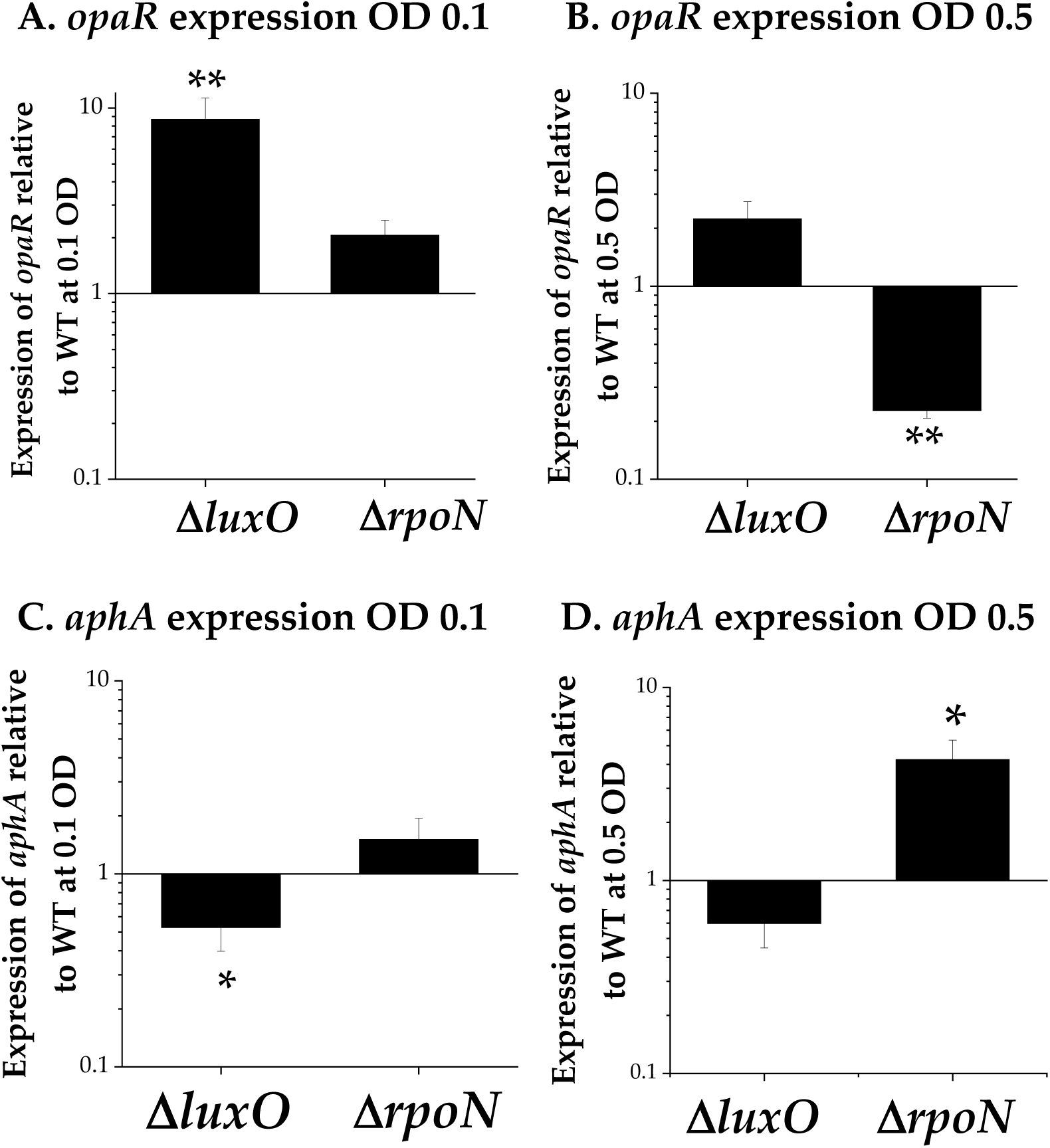
Quantitative real time PCR expression analysis of cells grown to 0.1 (A, C) or 0.5 OD (B, D) in LB media supplemented with 3% NaCl. Expression of *opaR* and *aphA* relative to WT RIMD2210633 and normalized to 16S housekeeping gene. Means and standard error of at least two biological replicates shown. Statistics calculated using a Student’s t-test. *, P-value <0.05; **, P-value <0.01.

### Expression analysis of *qrr1-qrr5* in Δ*luxO* and Δ*rpoN* mutants

Since *opaR* showed different levels of expression in the Δ*luxO* and Δ*rpoN* deletion mutants, we wanted to determine whether this was due to differences in *qrr* expression levels. We examined expression of all five *qrr* genes in cells grown to OD 0.1 and OD 0.5 and show that the expression of *qrr1, qrr2, qrr3* and *qrr5* was higher at OD 0.1 relative to OD 0.5 (**Fig. S2)**. Expression of *qrr4* was detected at OD 0.1 but was not detected at OD 0.5 in wild type. In addition, *qrr4* expression was not detected in either Δ*luxO* or Δ*rpoN* at OD 0.1 or 0.5. Next, we examined expression of the *qrr* genes at OD 0.1 in the Δ*luxO* mutant relative to wild type and showed *qrr1* expression was unchanged and there was significant downregulation of *qrr2, qrr3,* and *qrr5* (**Fig 4A**). Whereas at OD 0.5, their expression matched that of wild type (**Fig. 4B**). In the Δ*rpoN* mutant, expression of *qrr1, qrr3*, and *qrr5* matched that of the Δ*luxO* mutant (**Fig. 4C**), however, *qrr2* was upregulated at both OD 0.1 and OD 0.5 (**Fig. 4D**). To confirm that *qrr2* was differentially regulated between Δ*luxO* and Δ*rpoN*, the *qrr2* regulatory region was cloned into pRU1064 reporter vector upstream of a promoter-less *gfp* cassette (P*qrr2*-*gfp*). The specific fluorescence of P*qrr2*-*gfp* was examined in the wild type, Δ*luxO,* and Δ*rpoN* mutants and measured as a cumulative read-out of *qrr2* transcription (**Fig. 5A**). The level of specific fluorescence of P*qrr2-gfp* was reduced in the Δ*luxO* mutant relative to wild type, whereas in the Δ*rpoN* mutant, fluorescence was significantly increased (**Fig. 5A**). Next, we examined the *opaR* regulatory region cloned into pRU1064 reporter vector upstream of a promoter-less *gfp* cassette (P*opaR*-*gfp*) in wild type, a Δ*qrr2* single mutant and a Δ*qrr3,1,4,5* quadruple mutant with only *qrr2* present (**Fig. 5B**). In Δ*qrr2* compared to wild type, P*opaR*-*gfp* showed significantly increased fluorescence, whereas the quadruple *qrr* deletion mutant, with *qrr2* present, was similar to wild type (**Fig. 5B**). We predicted that deletion of *qrr2* in the Δ*rpoN* mutant background should restore *opaR* expression and CPS production. We constructed a Δ*rpoN/*Δ*qrr2* double mutant and examined *opaR* and *aphA* expression levels (**Fig. S3)**. Quantitative real time PCR assays showed that *opaR* was highly expressed in a Δ*rpoN/*Δ*qrr2* double mutant compared to wild type (**Fig. S3**).

**Figure 4:**
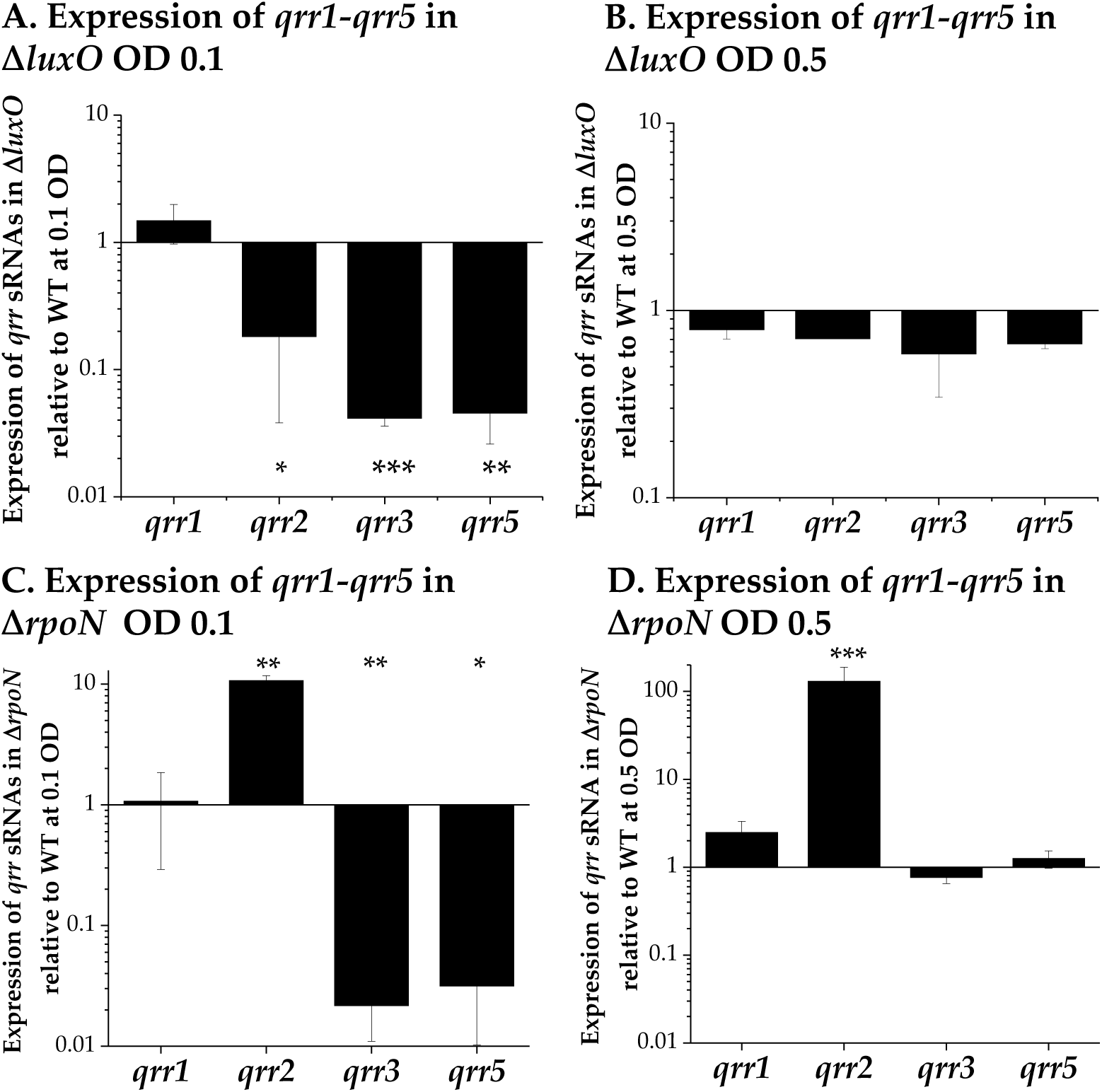
Quantitative real time PCR expression analysis of cells grown to 0.1 (**A, C**) or 0.5 OD (**B, D**) in LB media supplemented with 3% NaCl. Expression of *qrr1-5* relative to wild type RIMD2210633 and normalized to 16S housekeeping gene. Expression of *qrr4* not detected in mutant strains. Means and standard error of at least two biological replicates shown. Statistics calculated using a Student’s t-test. *, P-value <0.05; **, P-value <0.01; ***, P-value <0.001.

**Figure 5:**
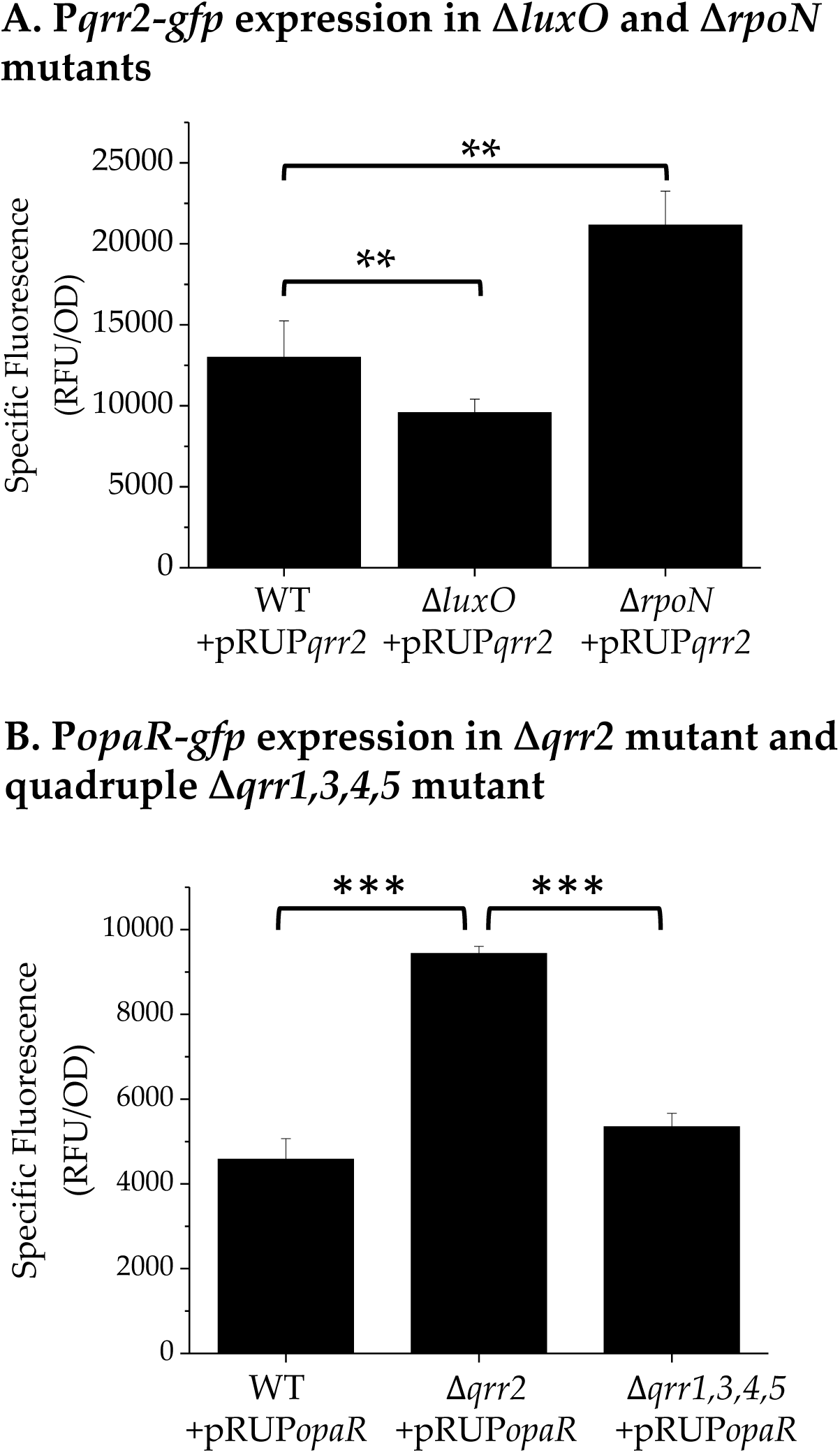
**A.** P*qrr2*-gfp reporter assay of *qrr2* in *luxO* and *rpoN* mutants. **B**. P*opaR*-gfp reporter assays in a single qrr2 deletion mutant and a quadruple mutant with only *qrr2* present. Cultures grown for 20 h in LB 3% NaCl. Means and standard error of at least three biological replicates shown. Statistics calculated using a one-way ANOVA and Tukey-Kramer *post-hoc* test. **, P-value <0.01

Examination of CPS formation showed that the Δ*rpoN/*Δ*qrr2* double mutant produced a rough colony morphology (**Fig. S4A**). Similarly, in biofilm assays, the Δ*rpoN* mutant produced a significantly reduced biofilm, whereas the Δ*rpoN/*Δ*qrr2* double mutant produced a biofilm similar to wild type (**Fig. S4B**). Overall, these data demonstrate that Qrr2 sRNA is present in the Δ*rpoN* deletion mutant and Qrr2 sRNA can function autonomously to control OpaR and QS phenotypes.

### Overlapping sigma-70 and sigma-54 promoters

The expression of *qrr2* in the Δ*rpoN* mutant background indicates that an additional sigma factor can initiate *qrr2* transcription. To examine this further, the regulatory regions of *qrr1* to *qrr5* in *V. parahaemolyticus* RIMD2210633 were aligned and, using bioinformatics tools, surveyed for the presence of promoter regions. Although the five Qrr sRNAs share homology, their regulatory regions are divergent with the exception of the sigma-54 canonical -24 (T**TGGC**A) and -12 (AAT**GCA**) promoter sites, with nucleotides in bold conserved amongst all five *qrr* regulatory regions (**Fig. S5**). In the regulatory region of *qrr2*, promoter analysis identified a housekeeping sigma-70 (RpoD) -35 (TTGAAA) and -10 (ATAATA) promoter (**Fig. 6A**). The putative sigma-70 promoter overlapped with the sigma-54 -24 and -12 promoter (**Fig. 6A**), and was absent from the regulatory regions of *qrr1, qrr3, qrr4,* and *qrr5* (**Fig. S5**). This suggested that *qrr2* can be transcribed by either sigma-54 or sigma-70 and could explain its expression in the absence of *rpoN*. To examine this further, we mutated three base-pairs of the putative sigma-70 -10 ATAATA site to ATACCC in the pRUP*qrr2* reporter vector (**Fig. 6A**). The mutagenized vector, pRUP*qrr2*-10CCC, was conjugated into wild type and Δ*rpoN* and specific fluorescence was determined. The Δ*rpoN* pRUP*qrr2*-10CCC strain showed significantly reduced fluorescence relative to Δ*rpoN* pRUP*qrr2*, indicating that this site is required for *qrr2* transcription in the absence of RpoN (**Fig. 6B**). The data suggests that *qrr2* can be transcribed by two sigma factors using dual overlapping promoters, suggesting a unique mode of regulation for Qrr2 sRNA. Comparisons of the *qrr2* regulatory region among Harveyi clade species *V. alginolyticus, V. campbellii, V. harveyi,* and *V. parahaemolyticus* showed that the sigma-70 promoter -10 region was highly conserved among these species (**Fig. S6**). Each of the five Qrr sRNAs also shared homology among these species (**Fig. S7**). The *qrr1* gene among all four species showed high homology clustering closely together on the phylogenetic tree, but were distantly related to the other four *qrr* genes. The *qrr3* and *qrr4* genes each clustered tightly together on the tree whereas *qrr2* and *qrr5* each showed divergence among the species (**Fig. S7**). Overall divergence in regulatory regions and gene sequence amongst the *qrr* genes likely suggests differences in how each *qrr* gene is regulated and differences in the target genes of each Qrr sRNA.

**Figure 6:**
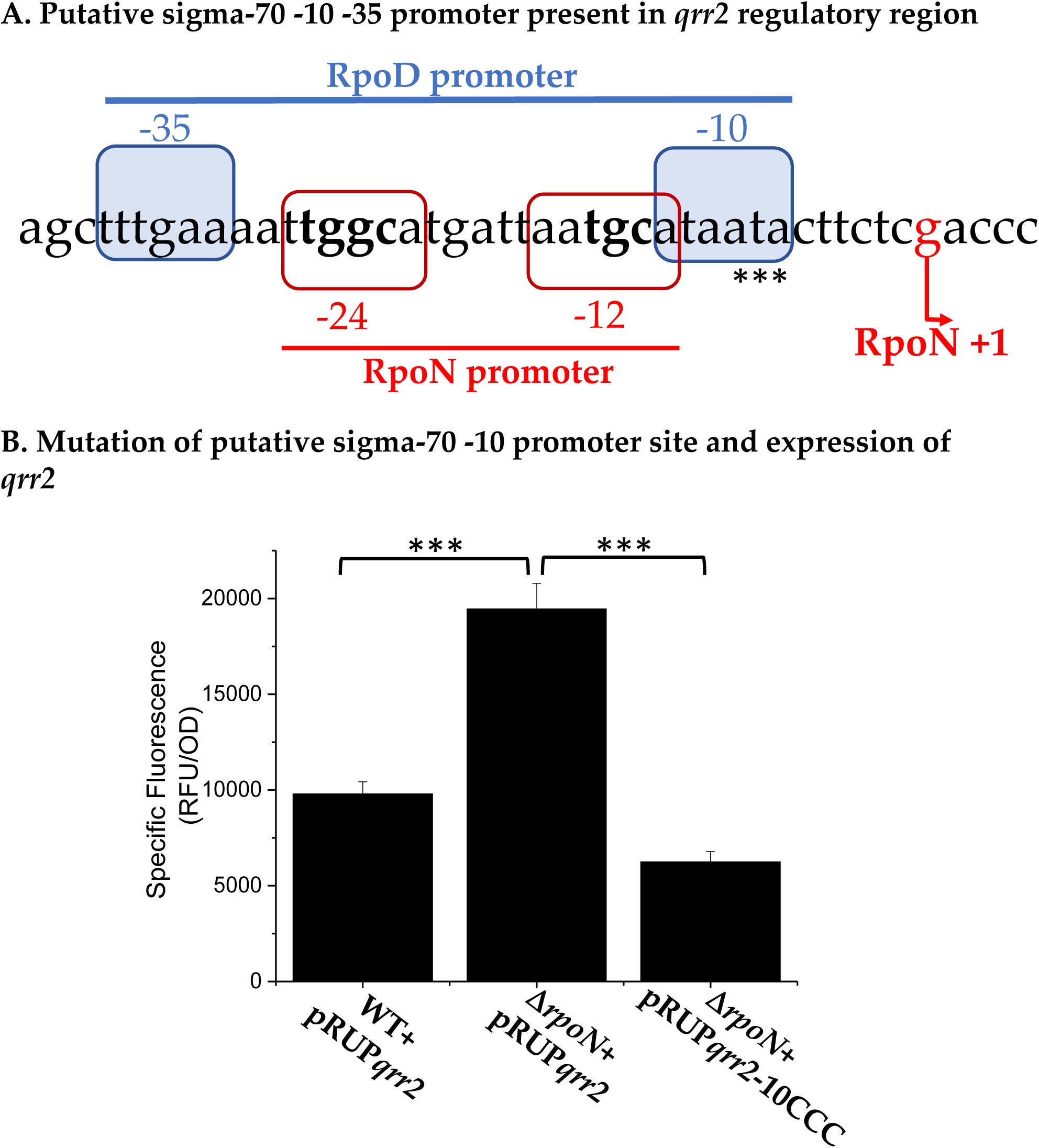
**A.** Analysis of *qrr2* regulatory region indicates overlapping sigma-54 and sigma-70 promoters. **B.** P*qrr2* GFP reporter assay of *qrr2* in Δ*rpoN* relative to wild type and mutated putative -10 RpoD binding site are indicated by asterisks. Means and standard error of three biological replicates shown. Statistics calculated using a one-way ANOVA and Tukey-Kramer *post-hoc* test. ***, P-value <0.001

### Qrr2 sRNA can function autonomously

Next, we determined whether Qrr2 sRNA has a distinct role in this species and whether any of the four other *qrr* genes can act independently. Using a *qrr1-qrr5* quintuple deletion mutant (Δ*qrr*-null) and five quadruple *qrr* deletion mutants, each containing a single *qrr*, we examined several QS phenotypes (**Fig. S8**). In swarming motility assays, the Δ*qrr*-null strain was swarming deficient, as swarming is negatively regulated by OpaR **(Fig. S8A)**. In addition, four quadruple mutants, Δ *qrr3,2,4,5;* Δ*qrr2,1,4,5;* Δ*qrr3,2,1,5;* and Δ*qrr3,2,1,4* were all swarming deficient indicating that Qrr1, Qrr3, Qrr4 and Qrr5 sRNAs cannot function independently to control this phenotype (**Fig. S8A)**. In swarming motility assays, the Δ*qrr3,1,4,5* mutant that contained only *qrr2*, behaved similar to wild type and was swarming proficient **(Fig. S8A)**. In swimming assays, the quad mutants that lacked *qrr2* produced similar results to the null mutant with defects in swimming (**Fig. S8B)**.

Whereas only Δ*qrr3,1,4,5* that contains only *qrr2* showed swimming motility similar to wild type (**Fig. S8B**). Additionally, in CPS assays the *qrr2* positive strain also showed a colony morphology similar to wild type (**Fig. S8C**). Analysis of a single *qrr2* deletion mutant indicates that it is not essential for CPS production or swarming and that the other *qrr* genes can function in the absence of *qrr2* (**Fig. S9**). In summary, these data demonstrate that only Qrr2 sRNA can function independently in *V. parahaemolyticus*.

### RpoN and Fis are not required for *qrr2* expression

In order to identify additional regulators of *qrr2* transcription, a DNA-affinity pull-down was performed. We used Δ*rpoN* cell lysate grown to OD 0.5 and P*qrr2* bait DNA. We identified a number of candidate regulators previously shown to bind to the *qrr* sRNA regulatory regions in *V. harveyi* (27, 34, 51) (**Fig. S10 and S11**). We decided to examine the nucleoid associated protein Fis further since it is known to be a positive regulator of *qrr* sRNA expression and binds to the *qrr* sRNA regulatory regions in *V. harveyi* (27). We identified three putative Fis binding sites in the *qrr2* regulatory region using virtual footprint analysis based off the E. coli Fis consensus sequence. A Fis binding site was located adjacent to the -35 promoter site, as well as two additional Fis binding sites, at 193-bp and 229-bp upstream of the *qrr2* transcriptional start site (**Fig. 7A**). To confirm these Fis binding sites, we constructed four DNA probes of the *qrr2* regulatory region to use in electrophoretic shift mobility assays (EMSAs) with purified Fis protein. DNA probe 1 encompassed the entire *qrr2* regulatory region and showed Fis binding in a concentration dependent manner via EMSA (**Fig 7B**). DNA probe 1A encompassing the single binding site showed binding in a concentration dependent manner, and similarly Fis bound to probe 1C which contained the two putative sites **(Fig. 7B**). Probe 1B, which did not have a putative Fis binding site, showed non-specific binding. Next, we examined expression of the P*qrr2*-*gfp* reporter in wild type, Δ*rpoN* and a Δ*rpoN/*Δ*fis* double mutant and confirmed that expression of P*qrr2-gfp* was upregulated in the Δ*rpoN* mutant but was even more highly upregulated in the Δ*rpoN/*Δ*fis* double mutant (**Fig. 7C**). These data indicate that both Fis and RpoN can act as repressors of *qrr2* in *V. parahaemolyticus*, and Fis likely plays a role in enhancing sigma-54 binding.

**Figure 7:**
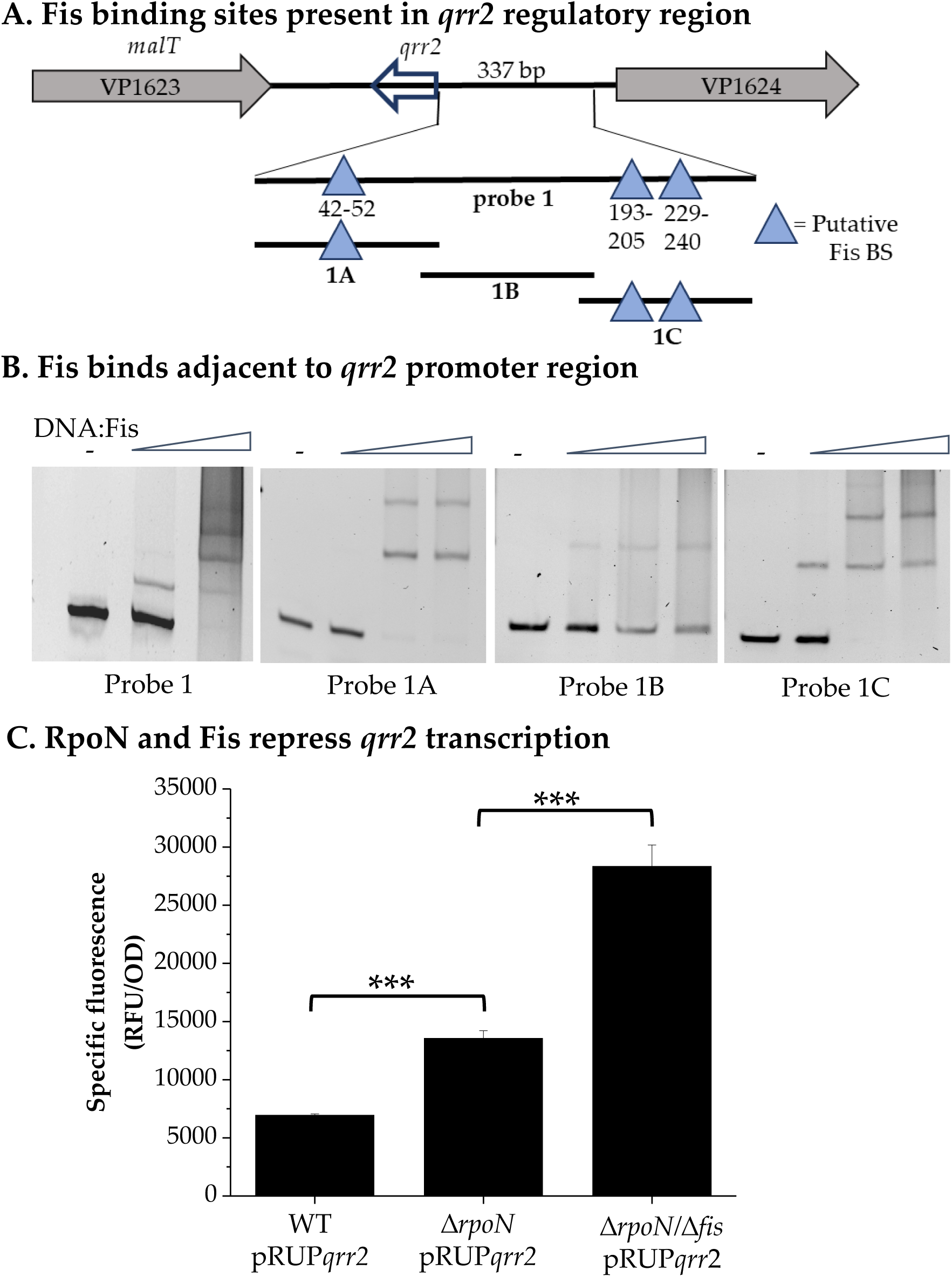
**A.** Regulatory region of *qrr2* depicted. Lines represent EMSA probes and blue triangles represent putative Fis binding sites using Virtual Footprint prediction software. Numbers indicate Fis binding site distance from *qrr2* transcriptional start site. **B.** Electrophoretic mobility shift assays of P*qrr2* with purified Fis protein using four *qrr2* regulatory region DNA probes **C.** pRUP*qrr2* reporter assays in Δ*rpoN* and Δ*rpoN/*Δ*fis* deletion mutants mutants relative to WT. Cultures grown for 20 h in LB 3% NaCl. Means and standard error of at least three biological replicates shown. Statistics calculated using a one-way ANOVA and Tukey-Kramer *post-hoc* test. ***, P-value <0.001.

## Discussion

In this study, we investigated the role of sigma-54, LuxO, and the five Qrr sRNAs in the *V. parahaemolyticus* QS pathway and showed that sigma-54, *qrr1, qrr3, qrr4,* and *qrr5* were not essential components. Our data demonstrated that in a Δ*rpoN* mutant, cells had a defect in CPS and biofilm formation, QS phenotypes that differed from the Δ*luxO* mutant. The data showed that Qrr2 is highly expressed in a Δ*rpoN* mutant and that Qrr2 can act autonomously to repress OpaR expression and QS phenotypes. In a Δ*rpoN/*Δ*qrr2* double mutant, *opaR* was de-repressed and CPS and biofilm formation were restored.

Bioinformatics analysis identified a putative -35 -10 promoter region within the *qrr2* regulatory region and mutagenesis of the -10 promoter sites resulted in repression of *qrr2*. Overall, the data indicate that *qrr2* can be expressed from two promoters, which is unique to *qrr2* in *V. parahaemolyticus,* but is likely true of related species. There have been other accounts of sigma-54-dependent genes showing increased transcription in the absence of *rpoN* (52, 53). In these cases, a putative sigma-70 promoter was present, suggesting a potential competition for promoter sites (52, 53). For example, in *E. coli, glmY* a coding sRNA contained overlapping sigma-54 and sigma-70 promoters, which were shown to allow for precise control of *glmY* expression within the cell (54). In our study, we identified a sigma-70 promoter that overlaps with the sigma-54 consensus promoter sequence of *qrr2*, suggesting that RpoN under differ growth conditions may block RpoD access. We propose that in the wild type background, *qrr2* is transcribed via LuxO activated RpoN, and in the Δ*luxO* mutant, *qrr2* is not transcribed because sigma-54 is in an inactive state bound to the *qrr2* promoter, physically blocking additional sigma factors from binding. However, in the absence of sigma-54, sigma-70 is able to bind to the *qrr2* regulatory region at a conserved -35 and -10 region to initiate transcription (**Fig. 8**). Fis is a global regulator that is known to enhance and inhibit transcription from promoter regions in many bacterial species (55-58). In *V. harveyi*, Fis has been shown to positively regulate *qrr* expression (27). Here, we show in DNA protein binding assays in *V. parahaemolyticus* that Fis binds adjacent to the -35 promoter site. We speculate that Fis functions to enhance RpoN promoter binding to maximize *qrr* expression. The data showed that in the absence of both RpoN and Fis, however, *qrr2* expression is significantly increased compared to the Δ*rpoN* mutant alone. Under these conditions additional binding sites within the *qrr2* regulatory region may be fully exposed, allowing sigma-70 full access for increased *qrr2* expression (**Fig. 8)**. A study in *V. alginolyticus* MVP01, a species closely related to *V. parahaemolyticus*, also showed differences between the Δ*luxO* and Δ*rpoN* mutant strains in their control of cell density dependent siderophore production. The Δ*luxO* mutant showed reduced siderophore production, which is negatively regulated by LuxR, and the Δ*rpoN* mutant showed increased production (59). Their data showed RpoN dependent and independent siderophore production. We speculate that this could be the result of expression by RpoD since *V. alginolyticus* has a -35 -10 promoter in the Qrr2 regulatory region (**Fig. S6**).

**Figure 8:**
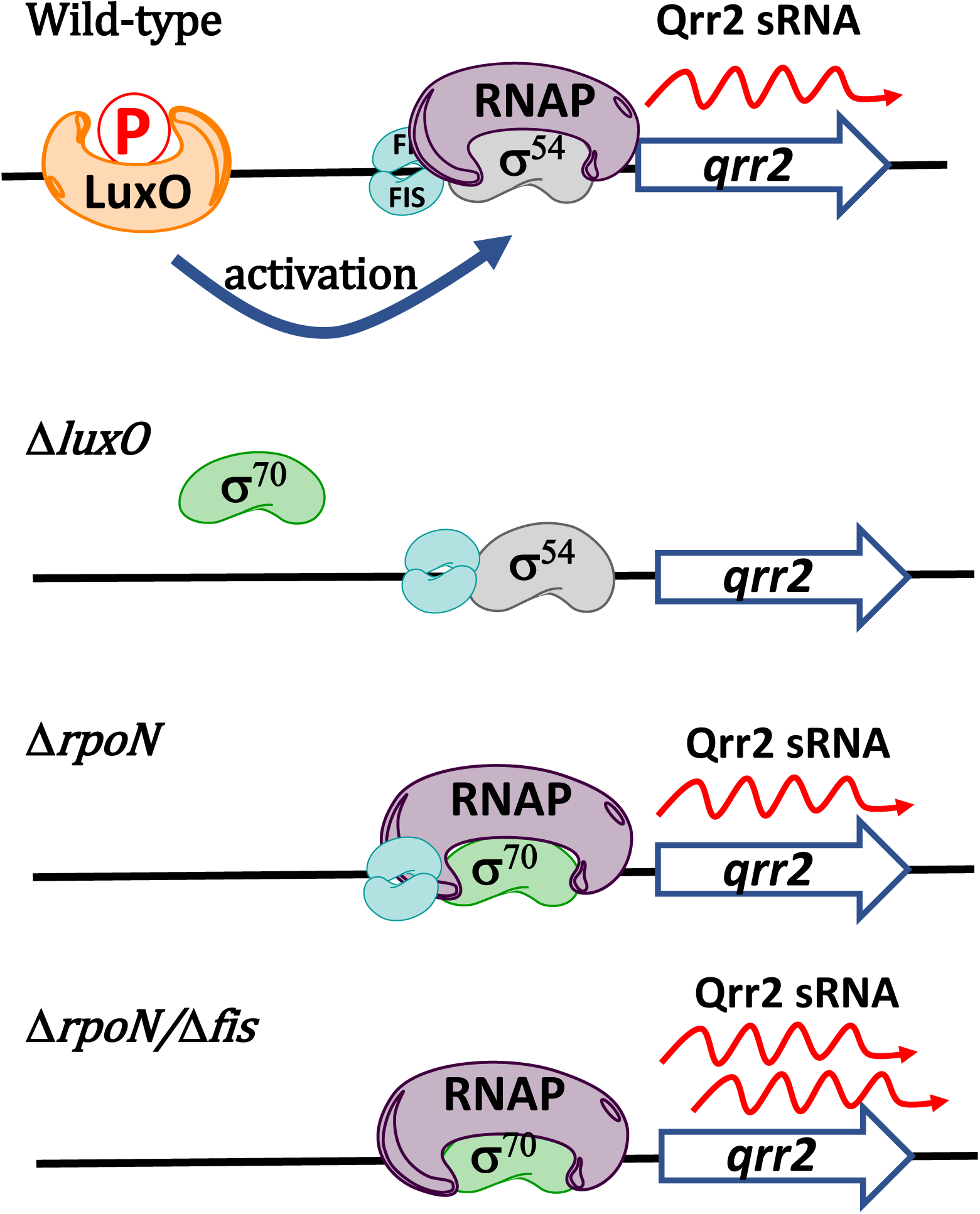
Model for *qrr2* transcription in the Δ*luxO* and Δ*rpoN* mutants. In the Δ*luxO* mutant, under certain conditions RpoN will be bound to the *qrr2* RpoN -24 -12 promoter region. RpoN bound at the promoter will be aided by Fis. This will prevent sigma-70 from binding. In the absence of RpoN (sigma-54), RpoD (sigma-70) can bind to the -35 -10 promoter region to initiate transcription. In the absence of Fis in the rpoN mutant transcription by RpoD is increased further as in the Δ*rpoN/*Δ*fis* mutant.

In *V. cholerae*, four Qrr sRNAs (Qrr1-Qrr4) are present that were shown to act redundantly to control bioluminescence, that is, any one of the Qrr sRNAs is sufficient to control HapR (LuxR homolog) (37). In their study, Lenz and colleagues showed that it was not until all four Qrrs were deleted in *V. cholerae*, that there is a difference in density-dependent bioluminescence (37). In *V. harveyi*, the five Qrrs were shown to act additively to control LuxR expression. Using bioluminescence assays and quadruple *qrr* mutants, it was determined that each Qrr has a different level of strength in repressing *luxR* translation (29). In our study in *V. parahaemolyticus*, the data showed that expression of *qrr4* was restricted to low cell density cells and *qrr4* expression had an absolute requirement for LuxO and RpoN. We demonstrated that Qrr2 sRNA is the only Qrr that can act autonomously to control QS gene expression, but is not essential, since a Δ*qrr2* mutant behaves like wild type (**Fig. S9**). Given that *qrr2* can be transcribed independent of RpoN, we propose that Qrr2 may have unique functions and/or targets in this species. We propose that *V. parahaemolyticus* can activate the transcription of *qrr2* via RpoN or RpoD to timely alter gene expression likely under different growth conditions.

## Materials and Methods

### Bacterial strains and media

In this study, the wild type (WT) strain is a streptomycin-resistant clinical isolate of *Vibrio parahaemolyticus* RIMD2210633 and all strains used are described in **Table 1**. All *V. parahaemolyticus* strains were grown in lysogeny broth (LB; Fisher Scientific, Fair Lawn, NJ) supplemented with 3% NaCl (LBS) (weight/volume). *E. coli* strains were grown in LB 1% NaCl. A diaminopimelic acid (DAP) auxotroph of *E. coli* β2155 λ*pir* was grown with 0.3 mM DAP in LB 1% NaCl. All strains were grown aerobically at 37°C. Antibiotics were used in the following concentrations: chloramphenicol (Cm), 12.5 μg/mL, streptomycin (Str), 200 μg/mL; and tetracycline (Tet), 1 μg/mL.

**Table 1.**
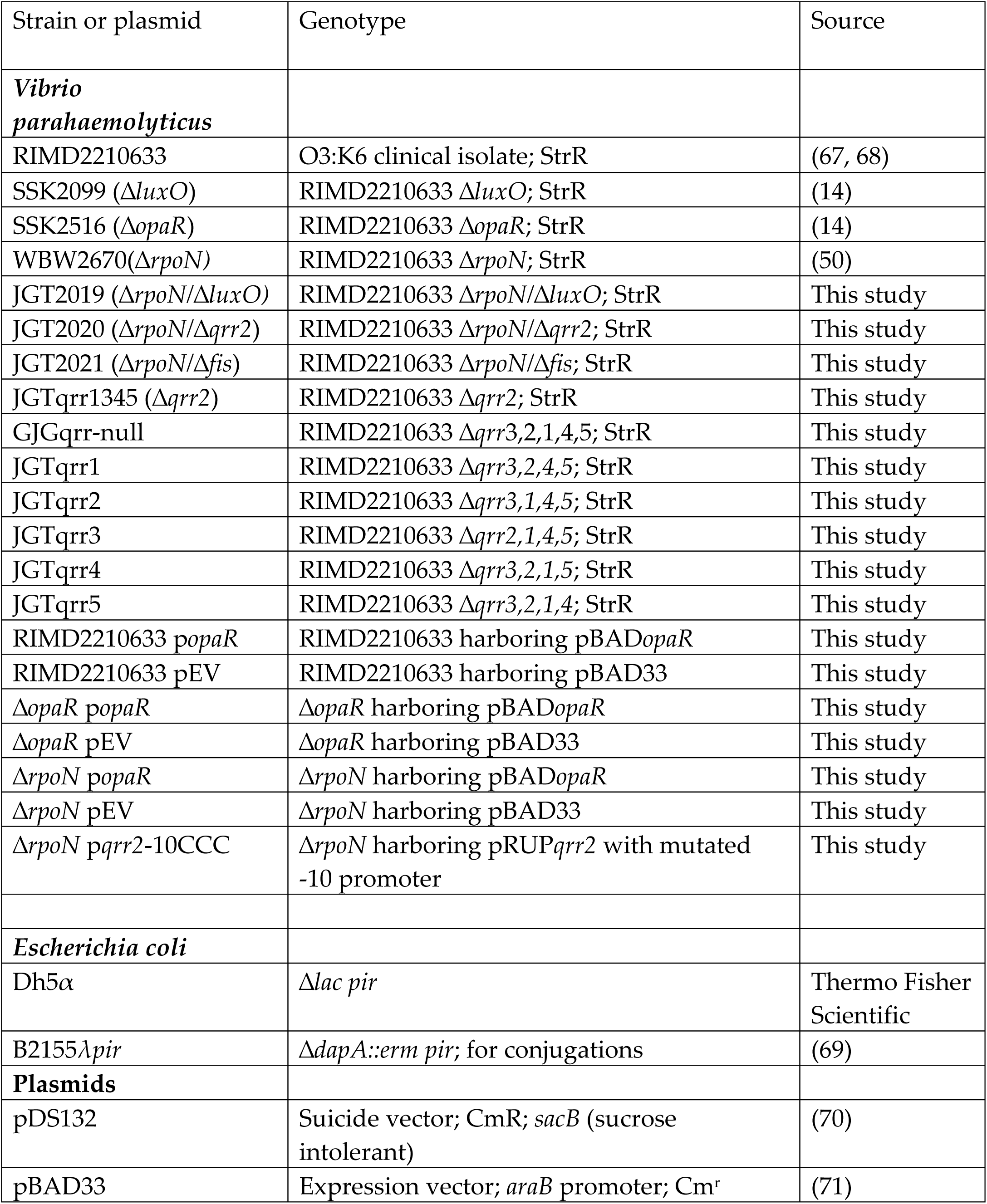
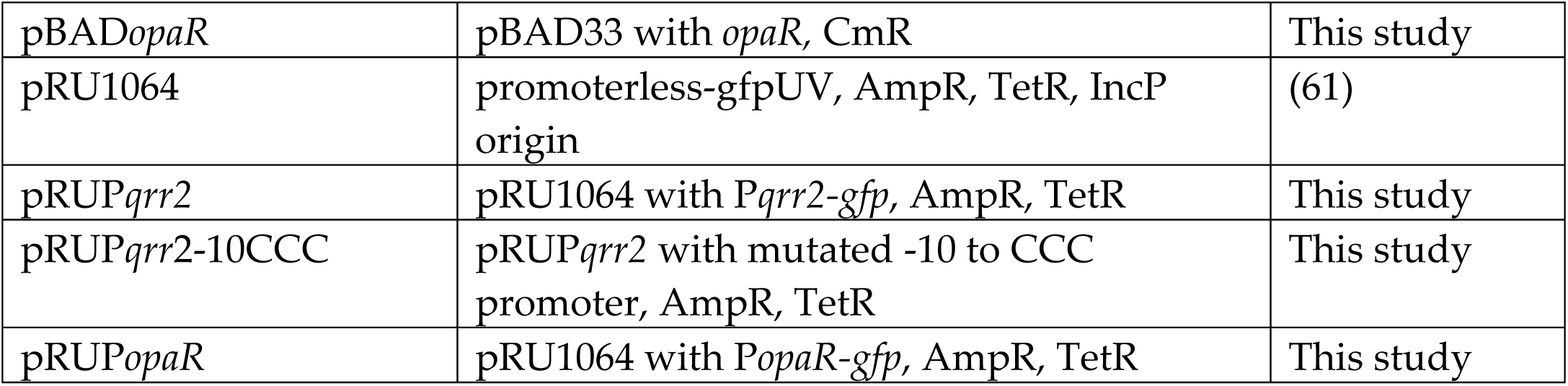
Strains and plasmids used in this study

### Construction of *V. parahaemolyticus* mutants

We created the double deletion mutants Δ*rpoN/*Δ*luxO* and Δ*rpoN/*Δ*fis* using mutant vectors pDSΔ*luxO* and pDSΔ*fis,* conjugated into the *V. parahaemolyticus* Δ*rpoN* mutant background. The Δ*qrr*-null mutant was constructed by creating truncated, non-functional copies of each *qrr* using SOE primer design, with primers listed in **Table 2** All truncated *qrr* products were cloned into pDS132 suicide vector, transformed into the *E. coli* β2155 λ*pir*, followed by conjugation and homologous recombination into the *V. parahaemolyticus* genome.

**Table 2.**
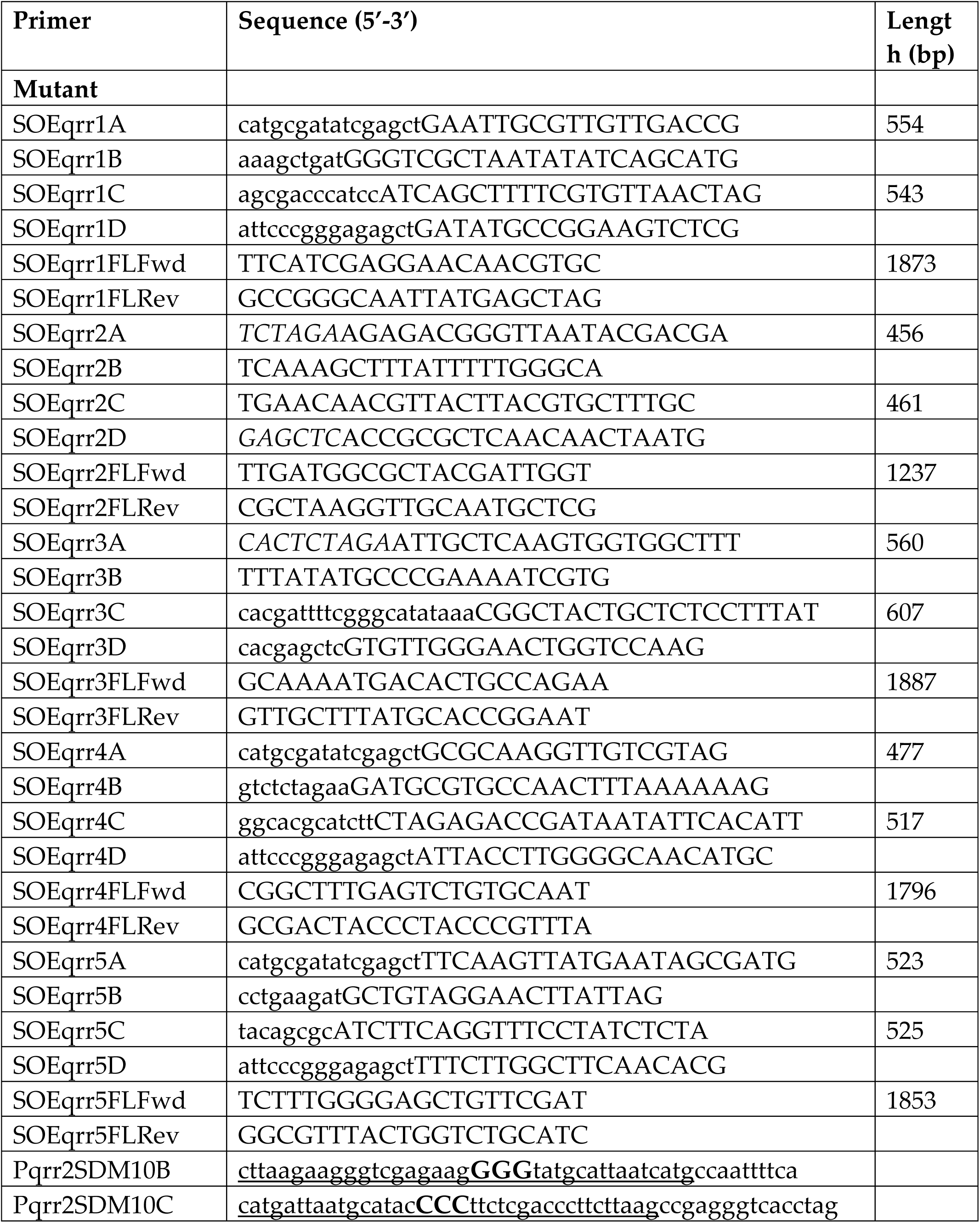
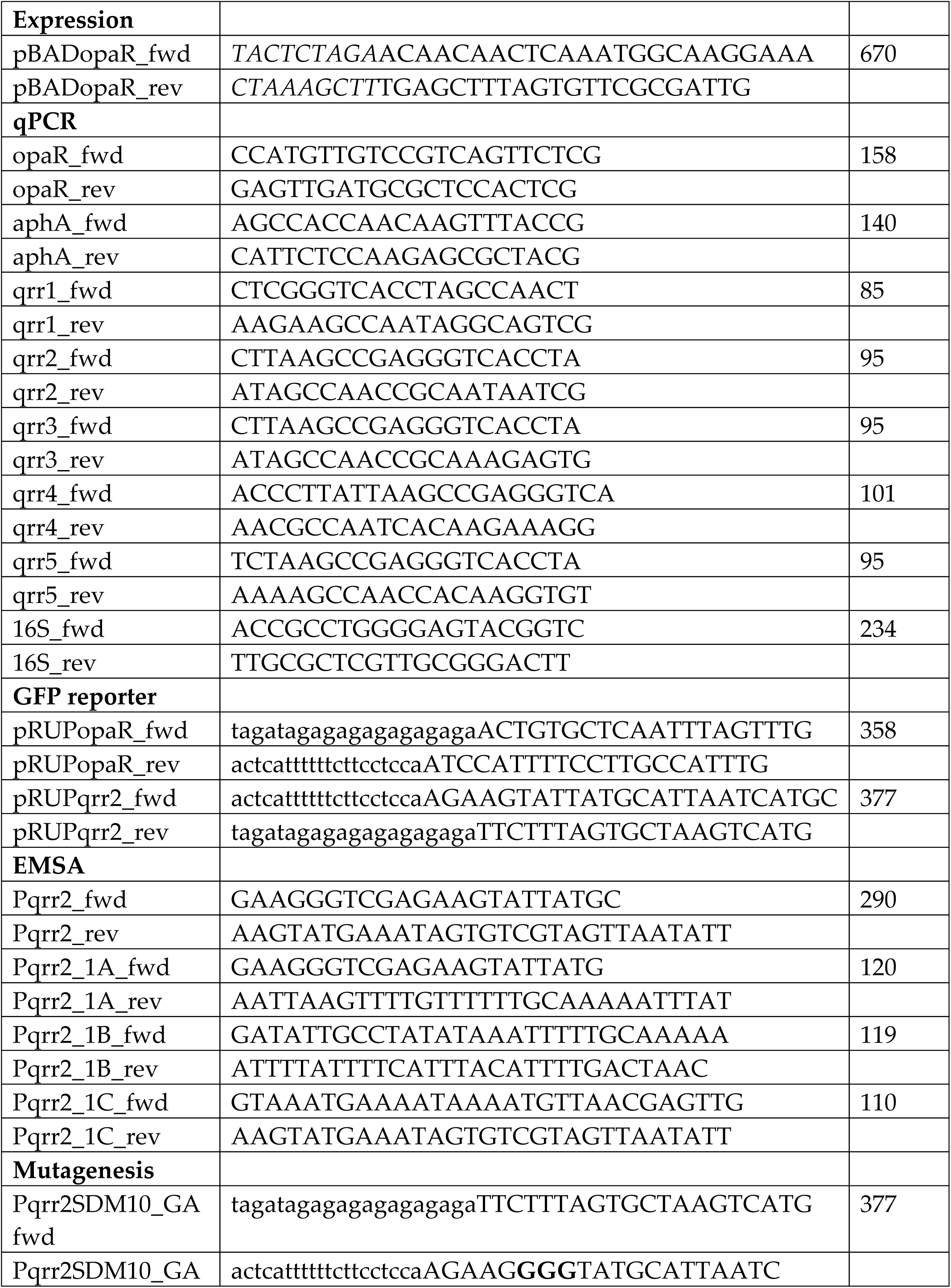
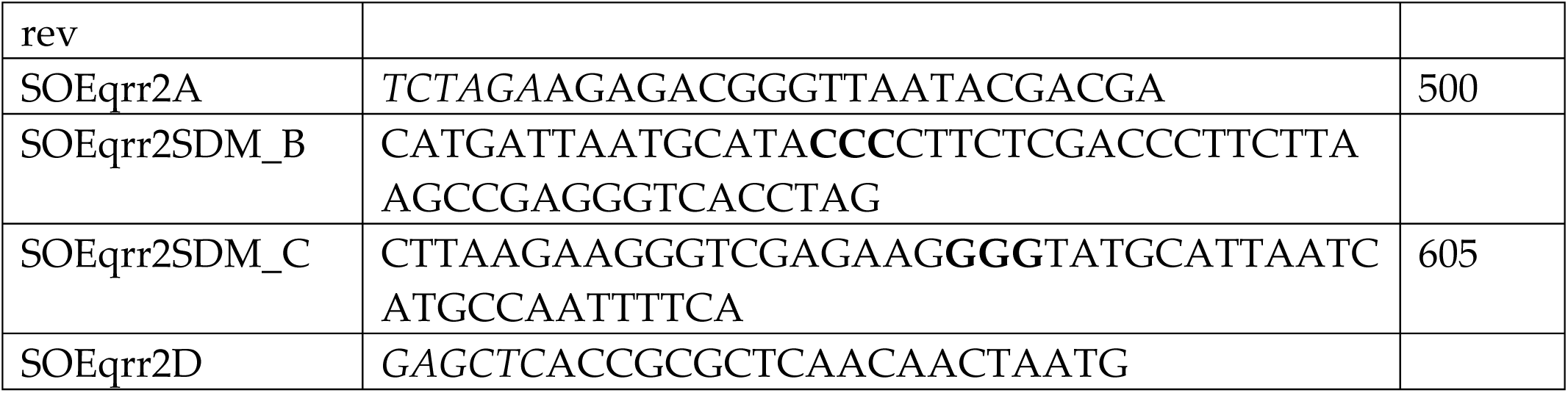
Primers used in this study

Positive single-cross over colonies were selected using Cm. To induce a double crossover event, a positive single-cross strain was grown overnight in the absence of Cm, leaving behind either the truncated *qrr* allele or the wild-type allele in the genome. The overnight culture was plated on sucrose plates for selection of normal versus soupy colony morphology, as the colonies still harboring the pDS132Δ*qrr* vector appear irregular due to the *sacB* gene. Colonies were screened via PCR for the truncated allele and sequenced to confirm deletion. The *qrr* null mutant was constructed by deleting qrr genes in the following order: *qrr3, qrr2, qrr1, qrr4, qrr5*. The quadrupleΔ*qrr3*/Δ*qrr2/*Δ*qrr4/*Δ*qrr5* mutant was constructed by re-introducing *qrr1* into the Δ*qrr*-null mutant, and similarly *qrr2* and *qrr3* were each separately cloned into the Δ*qrr*-null mutant to create their corresponding quad mutants. The Δ*qrr3*/Δ*qrr2/*Δ*qrr1/*Δ*qrr5* mutant was constructed by deleting *qrr5* in the Δ*qrr3*/Δ*qrr2/*Δ*qrr1* mutant background and Δ*qrr3*/Δ*qrr2/*Δ*qrr1/*Δ*qrr4* was constructed by knocking out *qrr4* in the Δ*qrr3*/Δ*qrr2/*Δ*qrr1* background. The Δ*qrr2* single mutant was constructed using the pDSΔ*qrr2* construct conjugated into the wild type background. The Δ*rpoN/*Δ*qrr2* mutant was constructed by conjugating the pDSΔ*qrr2* vector into the Δ*rpoN* background. All mutants were sequenced to confirm deletions or insertions, ensuring in-frame mutant strains.

### RNA isolation and real-time PCR

*Vibrio parahaemolyticus* wild type and mutants were grown overnight in LBS. Cells were washed twice with 1x phosphate-buffered saline (PBS) and diluted 1:50 into a fresh 5 mL culture of LBS. Cells were harvested at 0.1 OD and 0.5 OD and pelleted at 4°C. RNA was isolated from 4 mL of culture using the miRNAeasy Mini Kit (Qiagen, Hilden, Germany) and Qiazol lysis reagent. The concentration and purity of RNA was determined using a NanoDrop spectrophotometer (Thermo Scientific, Waltham, MA). RNA was treated with Turbo DNase (Invitrogen) and cDNA was synthesized using Superscript IV reverse transcriptase (Invitrogen) from 500 ng of RNA by priming with random hexamers. cDNA was diluted 1:10 for quantitative real-time PCR (qPCR) run on an Applied Biosystems QuantStudio™ 6 fast real-time PCR system (Applied Biosystems, Foster City, CA) using PowerUp SYBR green master mix (Life Technologies). qPCR primers used to amplify *opaR, aphA, qrr1, qrr2, qrr3, qrr4, qrr*5, and *16S* rRNA are listed in **Table 2** for reference. Cycle thresholds (C_T_) values were used to determine expression levels normalized to 16S rRNA levels. Expression was calculated relative to wild-type 16S rRNA using the ΔΔC_T_ method (60).

### Transcriptional GFP-reporter assay

The P*qrr2* reporter construct was created using the pRU1064 vector, which contains a promoter-less *gfp* cassette, as well as Tet and Amp resistance genes (61). Primers, listed in **Table 2**, were designed using NEBuilder online software to amplify the 337-bp regulatory region of *qrr2* from *V. parahaemolyticus* RIMD2210633 genomic DNA. The pRU1064 vector was purified, digested with Spe1, and ligated with the P*qrr2* fragment via Gibson assembly protocol (62). The plasmid was then transformed into β2155 λ*pir* and subsequently conjugated into wild-type and Δ*luxO*, Δ*rpoN,* and Δ*rpoN/*Δ*fis* mutants. Cultures were grown overnight in LBS with 1μg/mL Tet, washed twice with 1xPBS and then diluted 1:1000 into fresh LBS + Tet and grown for 20 hours at 37°C. Cultures were washed twice with 1xPBS and loaded into a black, clear-bottom 96-well plate. Final OD and GFP relative fluoresces were determined using a Tecan Spark microplate reader with Magellan software with excitation at 385 nm and emission at 509 nm (Tecan Systems, Inc., San Jose, CA). Specific fluorescence was calculated by dividing the relative fluorescence by the final OD. This experiment was performed in three biological replicates.

Splicing by overlap extension (SOE) primer design was used to construct a mutated (ATA-10CCC) RpoD promoter. We used the same SOE*qrr2*A and SOE*qrr2*D primers used to construct the Δ*qrr2* mutant in order to create a mutated *qrr2* regulatory region. In addition, SOE primers P*qrr2*SDMB and P*qrr2*SDMC (**Table 2**) have complementary overlapping sequences that amplify a mutated promoter, indicated in bold. Fragments AB and CD were then used as a template to amplify the AD fragment, containing a mutated RpoD -10 promoter. The AD fragment was then used as the template to create a fragment containing only the *qrr2* regulatory region (337-bp) using Gibson assembly primers P*qrr2*SDM_GAfwd and P*qrr2*SDM_GArev. This mutated regulatory region was then ligated with SpeI digested pRU1064 using Gibson assembly and confirm via sequencing.

### Capsule polysaccharide (CPS) formation assay

Capsule polysaccharide (CPS) formation assays were conducted as previously described (14, 45). In brief, single colonies of wild type and QS mutants were grown on heart infusion (HI) (Remel, Lenexa, KS) plates containing 1.5% agar, 2.5 mM CaCl_2_, and 0.25% Congo red dye for 48 h at 30°C. Each image is an example from at least three biological replicates. The pBAD33 expression vector was used to overexpress *opaR* in wild type and Δ*rpoN* backgrounds. The *opaR* coding region, plus 30-bp upstream to include the ribosomal binding site, were amplified from *V. parahaemolyticus* RIMD2210633 genome via Phusion High-Fidelity (HF) polymerase PCR (New England Biolabs). The amplified 670-bp *opaR* coding region and pBAD33 empty vector (pBADEV) were digested with XbaI and HindIII restriction enzymes prior to ligation and transformation into *E. coli* β2155. pBAD*opaR* and pBADEV were conjugated into wild type, Δ*rpoN,* and Δ*opaR,* and plated on Congo red plates to observe CPS formation. For strains containing pBAD, 0.1% (wt/vol) arabinose and 5 μg/mL of Cm were added to the media after autoclaving, to induce and maintain the plasmid, respectively.

### Biofilm assay

*Vibrio parahaemolyticus* cultures were grown overnight in LBS at 37°C with shaking. The overnight cultures were then used to inoculate a 96-well plate in a 1:50 dilution with LBS. After static incubation at 37°C for 24 h, the culture liquid was removed and the wells were washed with 1xPBS. Crystal violet (Electron Microscopy Sciences), at 0.1% w/v, was added to the wells and incubated for 30 min at room temperature. The crystal violet was removed, and wells were washed twice with 1xPBS. The adhered crystal violet was solubilized in DMSO for an optical density reading at 595nm (OD_595_).

### Motility assays

Swimming and swarming assays were performed as previously described (14, 50). To assess swimming, a pipette tip was used to pick a single colony and stab into the center of an LB plate containing 2% NaCl and 0.3% agar. Plates were incubated for 24 h at 37°C. Three biological replicates were performed, and the diameter of growth was measured for quantification. Swarming assays were conducted on HI plates containing 2% NaCl and 1.5% agar and incubated at 30°C for 48 h before imaging.

### DNA-affinity pull-down

A DNA-affinity pull-down was performed using previously described methods, with modifications as needed (63-65). Bait DNA primers were designed to amplify the regulatory region of *qrr2* (346-bp) with a biotin moiety added to the 5’ end. In addition, a negative control bait DNA (VPA1624 coding region, 342-bp) was amplified. Both bait DNA probes were amplified using Phusion HF polymerase (New England Biolabs) PCR and purified using ethanol extraction techniques (66). A 5 mL overnight culture of Δ*rpoN* grown in LB 3% NaCl was used to inoculate a fresh 100 mL culture of LB 3% NaCl grown at 37°C with aeration. The culture was pelleted at 0.5 OD at 4°C for 30 min and stored overnight at 80°C. The cell pellet was suspended in 1.5 mL of Fastbreak lysis buffer (Promega, Madison, WI) and sonicated to shear genomic DNA. The Δ*rpoN* lysate was pre-cleared with streptavidin DynaBeads (Thermo Scientific, Waltham, MA) to remove non-specific protein-bead interactions. Beads were incubated with 200 μL of probe DNA for 20 min, twice. The Δ*rpoN* lysate and sheared salmon sperm DNA (10μg/mL), as competitive DNA, were incubated with the beads twice, and washed. Protein candidates were eluted from the bait DNA-bead complex using elution buffers containing increasing concentrations of NaCl (100mM, 200mM, 300mM, 500mM, 750mM and 1M). 6X SDS was added to samples along with 1mM β-mercaptoethanol (BME) and then boiled at 95°C for 5 min. A total of 25 μL of each elution was run on 2 stain-free, 12% gels and visualized using the Pierce™ Silver Stain for Mass Spectrometry kit (Thermo Scientific, Waltham, MA). P*qrr2* bait and negative control bait were loaded next to each other in order of increasing NaCl concentrations. Bands present in the P*qrr2* bait lanes, but not in the negative control lanes were selected and cut from the gel. Each fragment was digested separately with trypsin using standard procedures and prepared for Mass Spectrophotometry 18C ZipTips (Fisher Scientific, Fair Lawn, NJ). Candidates were eluted in 10 μL twice, pooled, and dried again using SpeedVac. Dried samples were analyzed using the Thermo Q-Exactive Orbitrap and analyzed using Proteome Discoverer 1.4.

### Fis protein purification

Fis was purified using the method previously described (48). Briefly, primer pair FisFWDpMAL and FisREVpMAL was used to amplify *fis* (VP2885) from *V. parahaemolyticus* RIMD2210633. The *fis* gene was cloned into the pMAL-c5x expression vector fused to a 6X His tag maltose binding protein (MBP) separated by a tobacco etch virus (TEV) protease cleavage site. Expression of pMAL*fis* in *E. coli* BL21 (DE3) was induced with 0.5 mM IPTG once the culture reached 0.4 OD_595_ and grown overnight at room temperature. Cells were harvested, suspended in lysis buffer (50 mM NaPO4, 200 mM NaCl, and 20 mM imidazole buffer [pH 7.4]), and lysed using a microfluidizer. The lysed culture was run over an IMAC column using HisPur Ni-NTA resin, followed by additional washing steps. Mass spectrometry was performed to confirm Fis protein molecular weight and SDS-PAGE was conducted to determine its purity along with A260/280 ratio analysis using a Nano drop.

### Electrophoretic mobility shift assay for Fis

Purified Fis was used to conduct EMSAs using conditions previously described (48). Briefly, 30 ng of DNA probe was incubated with various concentrations of Fis (0 to 1.94 μM) in binding buffer (10 mM Tris, 150 mM KCL, 0.1 mM dithiothreitol, 0.1 mM EDTA, 5% PEG, pH7.4) for 20 min. The concentration of Fis was determined using Bradford reagent. A 6% native polyacrylamide gel was pre-run for 2 h at 4°C (200V) with 1x Tris-acetate-EDTA (TAE) buffer. The incubated DNA-protein samples were then loaded onto the gel (10 μL) and run for 2 h in the same conditions. The gel was stained in an ethidium bromide bath (0.5μg/mL) for 15 min before imaging. P*qrr2* was further divided into a smaller probe to determine specificity of Fis binding to P*qrr2*.

## Supplementary Figure Legends

**Figure S1:** Complementation analysis of Δ*rpoN* mutant with *opaR.* Expression vectors pBAD*opaR* and pBAD33 (empty vector) were conjugated into the *V. parahaemolyticus rpoN* mutant and spot inoculated on to congo red plates supplemented with 0.1% arabinose and 5 μg/ml chloramphenicol.

**Figure S2:** Gene expression analysis of cells grown to 0.1 OD and 0.5 OD in LB 3% NaCl. Expression of *qrr1* to *qrr5* at 0.1 OD relative to 0.5 OD, and normalized to 16S housekeeping gene. Means and standard error of at least two biological replicates shown. Expression of *qrr4* not detected at 0.5 OD. Statistics calculated using a Student’s t-test. *, P-value <0.05; **, P-value <0.01.

**Figure S3: A.** Quantitative real time PCR (qPCR) analysis of *opaR* and **B.** *aphA*. A. qPCR analysis in single D*rpoN* mutant and double D*rpoN*/D*qrr2* mutant. Two bio-reps in duplicate performed. Means and standard error of at least two biological replicates shown. Statistics calculated using a Student’s t-test. **, P-value <0.01

**Figure S4:** Phenotypic analysis of Δ*rpoN/*Δ*qrr2* double deletion mutant

**A.** CPS production in double mutant. **B.** Biofilm assay and quantification from cultures grown for 24 h stagnant and stained with crystal violet. Three bio-reps in duplicate performed. Statistics calculated using a one-way ANOVA and Tukey-Kramer *post-hoc* test. ***, P-value <0.001

**Figure S5.** The *V. parahaemolyticus* Qrr sRNAs regulatory and coding regions were aligned using T-COFFEE Multiple Sequence Aligner. The RpoN conserved binding site at -24 -12 and the transcriptional start site are labeled. The red lines indicate the RpoN promoter region with a highly conserved TGGC(-24) and TGC(-12). An asterisk indicates conserved nucleotides among all five *qrrs*. The blue box depicts the putative RpoD promoter of *qrr2*.

**Figure S6.** The *qrr2* gene sequence alignment from *V. harveyi* ATCC 33843*, V. campbellii* ATCC BAA-1116, *V. parahaemolyticus* RIMD2210633 *and V. alginolyticus* FDAARGOS_114 and the regulatory regions aligned using CLUSTALW. The sigma-54 conserved -24 -12 promoter binding sites are shown in blue boxes and the sigma-70 -35 and -10 promoter are shown in red boxes. The conserved nucleotides among the *qrr2* genes are shown by an asterisk. The data shows that the -10 promoter site is conserved between *V. parahaemolyticus* and *V. alginolyticus* with a 1-bp polymorphism in *V. harveyi*. Similarly, the -35 site is highly conserved between *V. parahaemolyticus* and *V. alginolyticus* but contains several polymorphisms in *V. harveyi*.

**Figure S7.** Phylogenetic tree of the *qrr* genes in *V. harveyi* ATCC 33843*, V. campbellii* ATCC BAA-1116, *V. parahaemolyticus* RIMD2210633 *and V. alginolyticus* FDAARGOS_114. The evolutionary history was inferred by using the Maximum Likelihood method and Jukes-Cantor model in MEGA X (1, 2). The tree with the highest log likelihood (-467.92) is shown. The percentage of trees in which the associated taxa clustered together is shown next to the branches. Initial tree(s) for the heuristic search were obtained automatically by applying Neighbor-Join and BioNJ algorithms to a matrix of pairwise distances estimated using the Maximum Composite Likelihood (MCL) approach, and then selecting the topology with superior log likelihood value. A discrete Gamma distribution was used to model evolutionary rate differences among sites (3 categories (+*G*, parameter = 0.2492)). The tree is drawn to scale, with branch lengths measured in the number of substitutions per site. This analysis involved 20 nucleotide sequences.

1. Jukes T.H. and Cantor C.R. (**1969**). Evolution of protein molecules. In Munro HN, editor, *Mammalian Protein Metabolism*, pp. 21-132, Academic Press, New York.

2. Kumar S., Stecher G., Li M., Knyaz C., and Tamura K. (**2018**). MEGA X: Molecular Evolutionary Genetics Analysis across computing platforms. *Molecular Biology and Evolution* **35**:1547-1549.

3. Felsenstein J. (**1985**). Confidence limits on phylogenies: An approach using the bootstrap. *Evolution* **39**:783-791.

**Figure S8:** Phenotypes of *qrr* deletion mutants. **A.** Swarming assay conducted on heart-infusion media incubated at 30°C for 48 h. **B.** Swimming motility assay conduced on semi-solid agar plates grown at 37°C for 24 h. Swimming plate quantification of three biological replicates. Statistics calculated using Student’s t*-*test relative to Wild-type. ***, P-value <0.001. **C.** CPS assays conducted of strains of interest. Colonies grown on Congo red plates for 48 h at 30°C prior to imaging.

**Figure S9.** Phenotype of qrr2 single deletion mutant. **A.** CPS production Strains of interest were inoculated on congo red plates and incubated for 48 hours at 30°C. Wild Type RIMD2210633 was used as a positive control and Δ*opaR* was used as a negative control. **B**. Swarming motility. Strains of interest were inoculated on swarming plates and incubated for 48 hours at 30°C.

**Figure S10:** DNA affinity pull-down of *qrr2* regulatory region (+) used to identify candidate regulators. Increasing concentrations of NaCl used to elute bound proteins. VP1624 coding region used as negative control bait DNA (-). Adjacent elutions used for comparison purposes. Boxes indicate chosen bands sent for Mass Spectrometry analysis. Boxes labeled N chosen to eliminate cross-over.

**Figure S11:** List of candidates identified in DNA affinity pull-down. Candidates divided into three categories, based on previously determined function. Fis is highlighted in red, as the target candidate in this study.

